# Common signatures of differential microRNA expression in Parkinson’s and Alzheimer’s disease brains

**DOI:** 10.1101/2022.01.31.478486

**Authors:** Valerija Dobricic, Marcel Schilling, Ildiko Farkas, Djordje O Gveric, Jessica Schulz, Lefkos Middleton, Steve Gentleman, Laura Parkkinen, Lars Bertram, Christina M. Lill

**Author notes:** **Corresponding Author:** Dr. Christina Lill, MD, MSc, Translational Epidemiology Group, LIGA, Ratzeburger Allee 160, Haus V50, 1st floor, Room 203.1, 23562 Lübeck, Germany. **Authors’ email addresses:** Valerija Dobricic, Marcel Schilling, Ildiko Farkas, Djordje O Gveric, Jessica Schulz, Lefkos Middleton, Steve Gentleman, Laura Parkkinen, Lars Bertram, Christina M. Lill.

## Abstract

**Background:** Dysregulation of microRNA (miRNA)-mediated gene expression has been implicated in the pathogenesis and course of many neurodegenerative diseases including Parkinson’s disease (PD). However, the functionally relevant miRNAs remain largely unknown. Previous meta-analyses on differential miRNA expression data in *post-mortem* PD brains have highlighted several miRNAs showing consistent and statistically significant effects. However, these meta-analyses were based on exceedingly small sample sizes.

**Methods:** In this study, we quantified the expression of the four most compelling PD candidate miRNAs from these meta-analyses in the superior temporal gyrus (STG) of one of the largest case-control *post-mortem* brain datasets available (261 samples), thereby quadruplicating previously investigated sample sizes. Furthermore, we probed for common differential miRNA expression signatures with Alzheimer’s disease (AD) by also analyzing these miRNAs in *post-mortem* STG of 190 AD patients and controls and by testing six top AD miRNAs in the PD brains.

**Results:** Of all ten analyzed miRNAs, PD candidate miRNA homo sapiens (hsa-) miR-132-3p showed evidence for differential expression in both PD (p=4.89E-06) and AD (p=3.20E-24), and AD miRNAs hsa-miR-132-5p (p=4.52E-06) and hsa-miR-129-5p (p=0.0379) showed evidence for differential expression in PD. Combining these novel data with previously published data substantially improved the statistical support (α=3.85E-03 using Bonferroni correction) of the corresponding meta-analyses clearly and compellingly implicating these miRNAs in both PD and AD. Furthermore, hsa-miR-132-3p/-5p (but not hsa-miR-129-5p) showed association with neuropathological Braak PD staging (p=3.51E-03/p=0.0117), suggesting that these miRNAs may play a role in α-synuclein aggregation beyond the early disease phase.

**Conclusions:** Our study represents the largest independent assessment of recently highlighted candidate brain miRNAs in PD and AD *post-mortem* brain samples, to date. Our results implicate hsa-miR-132-3p/-5p and hsa-miR-129-5p to be differentially expressed in both PD and AD brains, potentially pinpointing shared pathogenic mechanisms across these neurodegenerative diseases.

## Background

Parkinson disease (PD) is the most common movement disorder and second most common neurodegenerative disorder affecting 1 to 2% of the general population over the age of 60 years with increasing incidences in industrialized populations (1,2). Currently, there is no curative or preventive therapy available for PD, which is in part attributable to our lack of understanding its etiology. More than 90% of the disease is genetically complex, i.e., it is determined by a combination and likely interaction of multiple genetic, environmental, lifestyle and other intrinsic risk factors (3). While genome-wide association studies (GWAS) have identified ^~^90 independent genetic risk variants in PD (4) and several environmental and lifestyle variables have been described as being associated with PD (5), a substantial fraction of the disease variance remains unexplained. In this context - as for other neurodegenerative diseases - epigenetic mechanisms have been suggested to play a significant role in the molecular architecture and pathogenesis of PD. One proposed mechanism is the action of microRNAs (miRNAs) (reviewed in e.g., (3,6)). MiRNAs are small non-coding RNAs that bind to messenger RNAs (mRNAs) and thereby inhibit their translation into proteins (7). However, despite an increasing amount of data published on the possible role of miRNAs in PD pathogenesis, the interpretation of the individual findings has often been impeded by inconclusive or even discrepant results across studies. This can be attributed to heterogeneity across tissues used for the analysis, application of different methods and analysis protocols, and use of small sample sizes (8). The latter is of particular relevance in expression studies of *post-mortem* brain samples due to the paucity of available biomaterial. In order to identify miRNAs of potential relevance in PD, our group recently performed a systematic review, including meta-analyses on all published miRNA gene expression studies comparing *post-mortem* brains of PD patients with control individuals; four miRNAs showed significant evidence for differential miRNA expression changes in PD vs controls (8). Despite having merged all available relevant data in the field, the sample sizes for the meta-analyses still remained comparatively small (median n=88 derived from a median of 3 datasets, interquartile range [IQR] 87-98). Similar observations were made in other brain diseases, such as Alzheimer’s disease (AD), where we performed similar meta-analyses across all published differential miRNA expression studies (9) and which were also, even after combining all available data, still based on small sample sizes (median n= 42.5 derived from a median of 3 datasets [IQR 23-85; ref. (9)]).

In order to independently assess the role of the four most promising brain miRNAs in PD, we investigated their expression in 261 *post-mortem* brain samples including 214 PD patients and 47 controls, thereby effectively quadruplicating previously investigated sample sizes. Furthermore, we assessed common signatures of differential miRNA expression across neurodegenerative diseases by also investigating the same four miRNAs in *post-mortem* brain samples of 190 AD patients and controls, and by assessing expressional changes of six of the most promising AD miRNAs (9) in PD brain samples (**Figure 1**). Finally, our previous meta-analyses were updated with both the results from our current study and novel data from the literature published since our last data freeze in October 2018.

**Figure 1.**
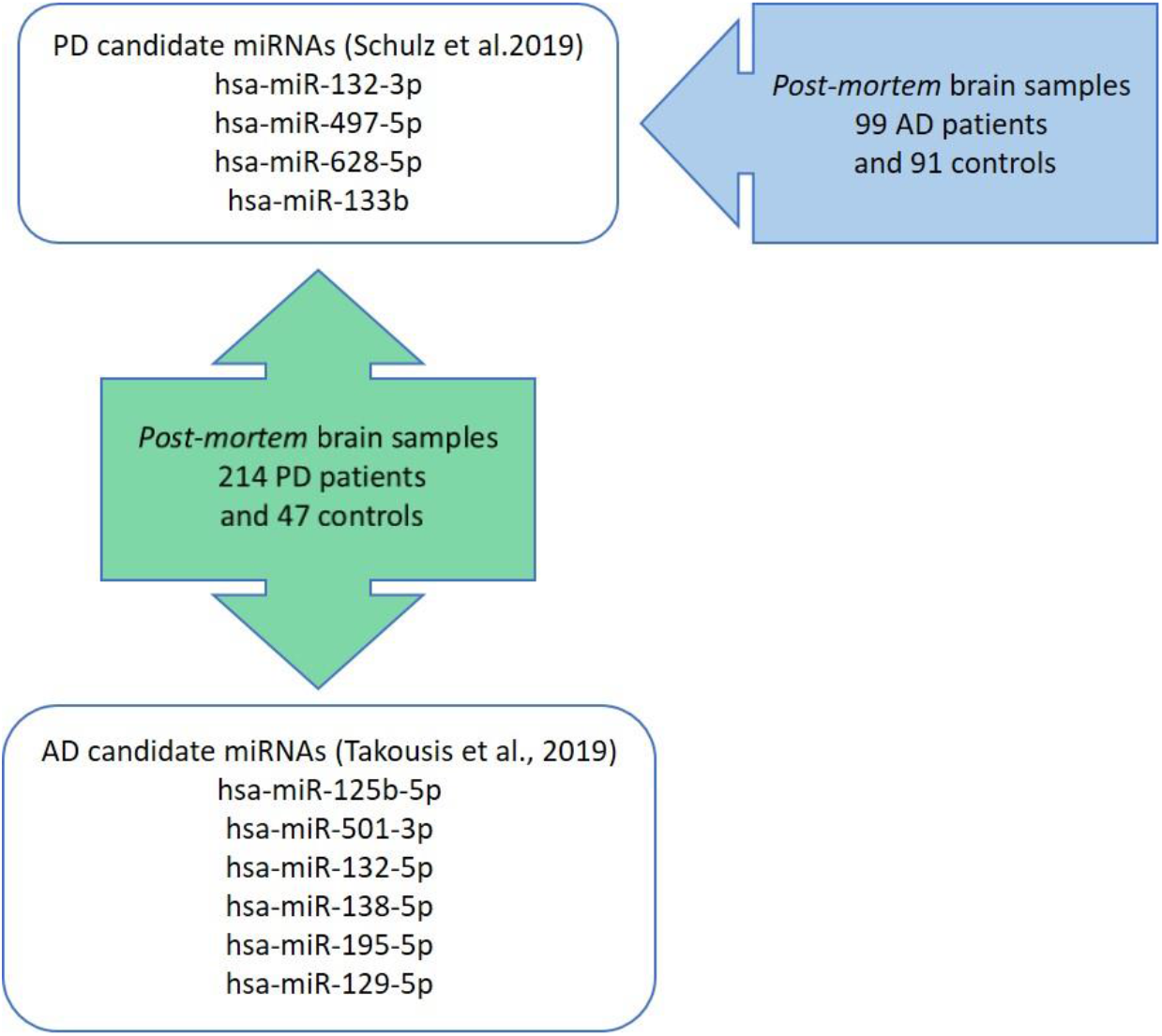
Overview of study design.

## Methods

### Brain samples

To investigate differential miRNA gene expression in PD and AD brains, we used superior temporal gyrus (STG) sections for this study, because this brain region has been identified as being selectively affected structurally and functionally at early stages in both PD (10) and AD (e.g., ref. (11)). For PD, snap-frozen STG sections (Brodmann area BA21) of 235 deceased patients with a clinical and neuropathologically confirmed diagnosis of PD and 47 controls were provided by the Parkinson’s UK Brain Bank at Imperial College London (ICL). Staging of α-synuclein and tau pathology was conducted according to the BrainNet Europe Consortium criteria (12,13). Presence and distribution of α-synuclein and tau deposits were assessed using immunohistochemistry, with antibodies specific for a-synuclein (mouse anti-α-synuclein, BD Transduction Laboratories, Franklin Lakes, New Jersey) and hyperphosphorylated tau (AT8, Invitrogen, Waltham, Massachusetts). For AD, we used snap-frozen STG sections (Brodmann area BA21) of 99 AD patients and 91 elderly control individuals provided by the Oxford Brain Bank (for a detailed description of the samples used here, see (14)). In brief, AD patients and controls were part of the longitudinal, prospective Oxford Project to Investigate Memory and Aging (OPTIMA) and underwent a standard battery of clinical and neuropsychological tests (15). The pathological diagnosis of AD was made using the CERAD/NIH criteria and Braak AD staging (16–18). All participants had given prior written informed consent for the brain donation. The tissue bank activities of the Parkinson’s UK Brain Bank were approved by the Ethics Committee for Wales (ref 18/WA/0238), and the Oxford Brain Bank activities were approved by the Ethics Committee of the University of Oxford (ref 15/SC/0639).

### MiRNA extraction and qPCR analysis

Total RNA (including miRNAs) was extracted from brain tissue using the mirVana isolation kit (Thermo Fisher Scientific, Waltham, Massachusetts) following the manufacturer’s instructions. For each extraction we used approximately 25mg of snap-frozen tissue. In order to minimize potential batch effects, patients and controls were randomly mixed before each extraction procedure. Residual DNA in RNA samples was removed using DNase (TURBO DNA-free kit, Thermo Fisher Scientific). RNA quantity was assessed using NanoDrop 2000 (Thermo Fisher Scientific), and RNA quality was assessed by determining RNA integrity numbers (RINs) using the Agilent 2100 Bioanalyzer system with the Agilent RNA 6000 Nano Chip Kit (Agilent Technologies, Santa Clara, California).

Reverse transcription reactions were performed on 10 ng of total RNA using the TaqMan Advanced miRNA cDNA Synthesis Kit (Thermo Fisher Scientific), as per manufacturer’s instructions. In the PD case-control brain samples we quantified a total of ten miRNAs: these included all four miRNAs showing significant differential expression in brain tissue of PD patients and controls in our recent meta-analysis (8), i.e., hsa-miR-132-3p (MIMAT0000426), hsa-miR-497-5p (MIMAT0002820), hsa-miR-628-5p (MIMAT0004809), and hsa-miR-133b (MIMAT0000770). In addition, we quantified the six most compelling miRNAs from our recent meta-analyses of differential miRNA expression studies in AD(9). These six miRNAs were among the top 10 most significantly associated miRNAs in AD brains in that study (9) and, in addition, showed little between-study heterogeneity (i.e., ≥80% of the datasets showed the same direction of effect): hsa-miR-125b-5p (MIMAT0000423), hsa-miR-501-3p (MIMAT0004774), hsa-miR-132-5p (MIMAT0004594), hsa-miR-138-5p (MIMAT0000430), hsa-miR-195-5p (MIMAT0000461), and hsa-miR-129-5p (MIMAT0000242). Furthermore, in the AD case-control brain samples, we assessed the expression of the four PD miRNAs from our previous study (8) (listed above). As endogenous controls, alongside all samples, we used hsa-miR-423-5p (suggested as endogenous control by ThermoFisher, Inc) for PD, and hsa-miR-423-5p and hsa-let-7b-5p for AD. All assays were run in a pre-spotted 384-well format on a QuantStudio-12K-Flex system (Thermo Fisher Scientific). Samples were measured in triplicate. Raw data analysis was performed using the ExpressionSuite Software v1.2 (Thermo Fisher Scientific). Samples for which either the endogenous control assay (n = 10) or ≥ 4 target miRNA assays failed (n = 11) were excluded from all subsequent analyses. In addition, for the samples passing sample QC (214 PD patients, 47 controls; 99 AD patients, 91 controls) individual assays with differences in Ct value (ΔCt) > 0.5 across the triplicate measurements were excluded. The exact number of samples for each assay included in all subsequent statistical analyses is given in **Table 1**.

**Table 1.** Differential expression analysis of PD candidate miRNAs in brains of PD and AD patients and controls

| PD candidate<br>miRNA | PD |  |  |  | AD |  |  |  |
| --- | --- | --- | --- | --- | --- | --- | --- | --- |
| | Relative quantity<br>(95% CI) | Effect size<br>(±SE) | P-value | N (PD cases,<br>controls) | Relative quantity<br>(95% CI) | Effect size<br>(±SE) | P-value | N (AD cases,<br>controls) |
| hsa-miR-132-3p | 0.567 (0.507, 0.635) | -0.708 (±0.151) | **4.89E-06** | 253 (211, 42) | 0.282 (0.248, 0.320) | -1.21 (±0.102) | **3.23E-24** | 190 (99, 91) |
| hsa-miR-133b | 0.999 (0.935, 1.067) | -0.0282 (±0.169) | 0.868 | 248 (204, 44) | 0.927 (0.837, 1.026) | 0.0879 (±0.168) | 0.602 | 186 (96, 90) |
| hsa-miR-497-5p | 1.050 (0.994, 1.110) | 0.0556 (±0.159) | 0.726 | 251 (206, 45) | 0.998 (0.940, 1.060) | 0.239 (±0.163) | 0.143 | 190 (99, 91) |
| hsa-miR-628-5p | 0.965 (0.882, 1.055) | -0.0902 (±0.153) | 0.556 | 246 (203, 43) | 0.769 (0.688, 0.858) | -0.226 (±0.151) | 0.137 | 189 (98, 91) |

**Legend.** This table displays the statistical results of the differential gene expression analyses of Parkinson's disease (PD) candidate miRNAs in *post-mortem* brain samples of 214 PD patients and 47 controls and of 99 Alzheimer's (AD) patients and 91 controls, respectively. P-values displayed in bold highlight nominally significant ( $\alpha=0.05$ ) differential expression results. SE = standard error; N = number

### Statistical analysis

Comparisons of age, *post-mortem* intervals (PMI), RIN values and RNA absorbances between patients and control samples were performed by Welch’s t-test, and, in case of non-normality, by Wilcoxon rank sum test, and the comparison of sex distributions was compared by the chi-squared test using R (https://www.R-project.org/). Differential gene expression analysis was performed by (independently) fitting a (Gaussian) generalized linear model (GLM) to predict each candidate miRNA expression (measured by qPCR ΔCt; scaled and centered) from disease status (PD/AD case vs. control) and potential confounding factors ( RIN, age at death, and PMI; scaled and centered; as well as sex). Significance was assessed by testing against the null hypothesis that the disease status does not contribute (i.e. zero-weight) to the model (two-sided z-test). The type-1-error rate was set to α = 0.05 for this candidate-based validation approach (see below for our conservative multiple testing correction of the corresponding meta-analyses).

Furthermore, we utilized PD cases to train GLMs predicting gene expression of the top miRNAs hsa-miR-132-3p/-5p and hsa-miR-129-5p based on α-synuclein and tau Braak staging (see above) as separate outcomes. To this end, the corresponding binary (case vs. control) variable was replaced by a continuous (scaled and centered) variable. All other analysis steps and covariate adjustments were identical to the case vs. control analyses. While we performed six tests in this arm of the study (testing α-synuclein and tau Braak staging for three miRNAs each), we note that levels for hsa-miR-132-3p and hsa-miR-132-5p are highly correlated (Pearson’s r = 0.93, p = 2.2E-16), and not independent. Thus, the type-1 error rate for this arm of the study was set to α = 0.05 / 4 = 0.0125 using Bonferroni correction for 4 independent tests for the neuropathological assessment.

### Literature search and meta-analyses

We assessed the overall evidence for differential expression of the ten PD and AD candidate miRNAs in PD brains by updating our earlier meta-analyses (8). To this end, we included the data on these ten miRNAs reported in the previous study (8), the molecular data generated in our current study and other recently published data, following the data freeze of our previous study (8). For the literature update, we applied the exact same methodology as described previously (8) and included all studies published until December 1^st^, 2021. Similarly, the meta-analyses performed in AD for the four PD candidate miRNAs (9) were updated using the same approach, i.e., combining the data previously reported by Takousis et al. (9), our newly generated data in the AD case-control dataset, and data published since the original data freeze (until December 1^st^, 2021).

To correct for multiple testing across meta-analyses we used Bonferroni’s method and adjusted for 13 independent tests: 10 miRNAs were tested for PD (nine of which were independent, see above), and 4 uncorrelated miRNAs were tested for AD resulting in a two-sided study-wide α = 0.05 / 13 = 3.85E-03 for the meta-analysis results. In addition, we pre-defined the requirement that a miRNA was to be considered as associated with PD and AD, respectively, only if it passed the study-wide type-1 error rate of a two-sided α = 3.85E-03 in the meta-analyses *and* if statistical significance improved in the meta-analysis combining previously published and newly generated data in comparison to the results described in Schulz et al., 2019 (8).

## Results

### Demographics and RNA quality assessments

The effective dataset analyzed in this study comprised *post-mortem* STG brain samples of 214 patients and 47 controls for PD (**Additional file 1, Table S1**), and of 99 patients and 91 controls for AD (**Additional file 2, Table S2**). The age at death, sex distribution, and *post-mortem* interval until collection of brain samplesdid not show significant differencesbetween PD patients and controls. For AD, there was a nominally significant difference in the age at death distribution and significant associations of RIN and RNA absorbance values with disease status (**Additional file 2, Table S2**). However, comparison of raw expression data in the AD dataset showed that the distribution of Ct values was similar for samples with lower (RIN < 5) vs higher RIN values (RIN ≥ 5) suggesting that sample quality has not substantially impacted our results (**Additional file 3, Figure S1**). To adjust for potential residual confounding due to these variables we included them as covariates in the linear regression model for both PD and AD (see below).

### Differential miRNA expression analysis in STG

Quantification of the four PD candidate miRNAs confirmed that hsa-miR-132-3p was strongly and significantly downregulated in our large independent collection of PD brain samples compared to controls (p = 4.89E-06; **Table 1, Figure 2a**). The other three miRNAs (hsa-miR-133b, miRNAs hsa-miR-497-5p, and hsa-miR-628-5p) did not show evidence for differential expression in PD (**Table 1, Figure 2a**). Assessment of these four PD candidate miRNAs in our AD dataset revealed that hsa-miR-132-3p was strongly downregulated not only in PD, but also in AD (p = 3.20E-24; **Table 1, Figure 2b**). The remaining three PD candidate miRNAs did not show significant differential expression in AD, similar to our observation in PD (**Table 1, Figure 2b**).

**Figure 2.**
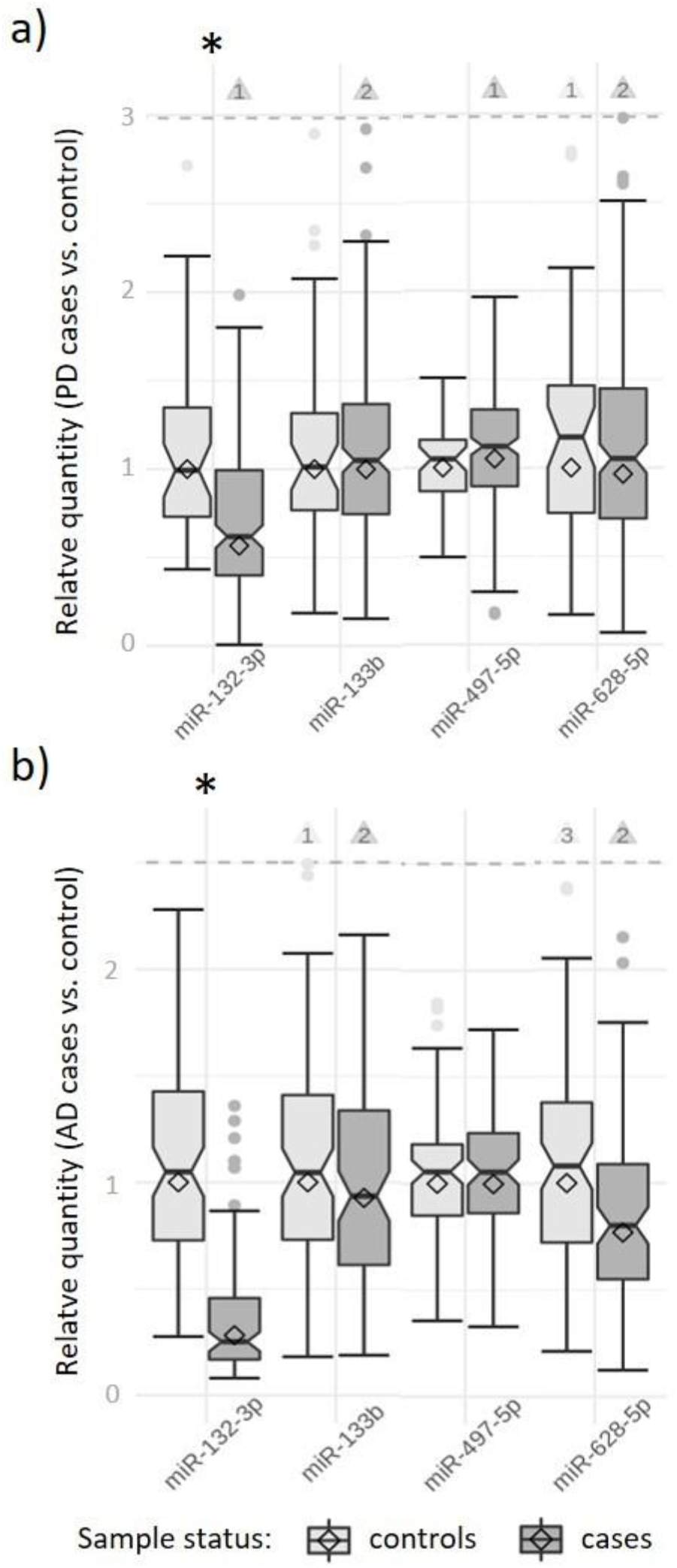
Expression levels of PD candidate miRNAs in brain samples of PD and AD patients versus controls. <u>Legend</u>. This box and whisker plot displays the relative quantity of Parkinson’s disease (PD) candidate miRNAs in *post-mortem* brain samples of the supratemporal gyrus of 214 PD patients vs. 47 controls (Figure 2a) and 99 AD patients vs. 91 controls (Figure 2b). The relative quantity of miRNA expression was calculated using the ΔΔCt method; diamonds represent the mean expression (cases relative to controls) based on the ΔΔCt method (relative quantity = 2(^-(dCt cases - dCt controls))^). Horizontal lines represent median values the corresponding sample-specific values (individual dCt values normalized to the mean of the control samples), boxes represent interquartile ranges, and whiskers extend to the maximum observed value within 1.5x the interquartile range; values outside this range but below the dashed line are depicted as dots. Outliers exceeding the dashed line are not shown (for scaling purposes) but counted and indicated by the numbers in the triangles. The box notches indicate the corresponding 95% confidence intervals. * = nominally statistically significant difference (α=0.05).

To further investigate cross-disease miRNA expression changes, we also quantified the six previously described AD candidate miRNAs in our PD dataset. These analyses revealed that hsa-miR-132-5p was significantly downregulated in PD brains compared to controls (p = 4.52E-06, **Table 2, Figure 3**). Further analyses showed that this miRNA was highly correlated with hsa-miRNA-132-3p in our data (Pearson’s r = 0.93, p = 2.2E-16, also see Methods). In addition, the AD candidate miRNA miR129-5p showed evidence for significant downregulation in PD (p = 0.0379). None of the other four AD candidate miRNAs (hsa-miR-125b-5p, hsa-miR-138-5p, hsa-miR-195-5p, and hsa-miR-501-3p) showed statistically significant differential expression in PD.

**Table 2.**
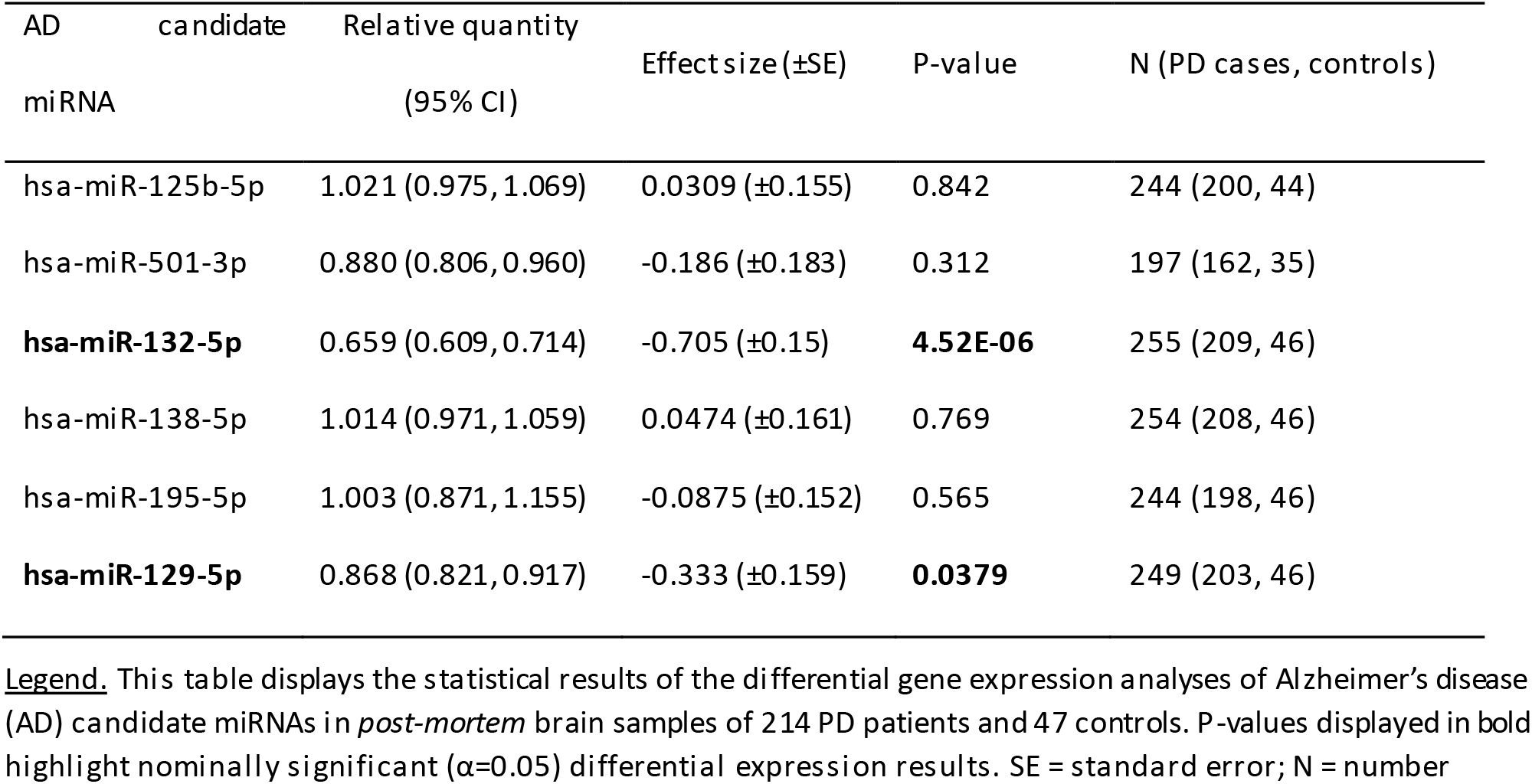
Differential expression analysis of AD candidate miRNAs in brain samples of PD patients and controls

**Figure 3.**
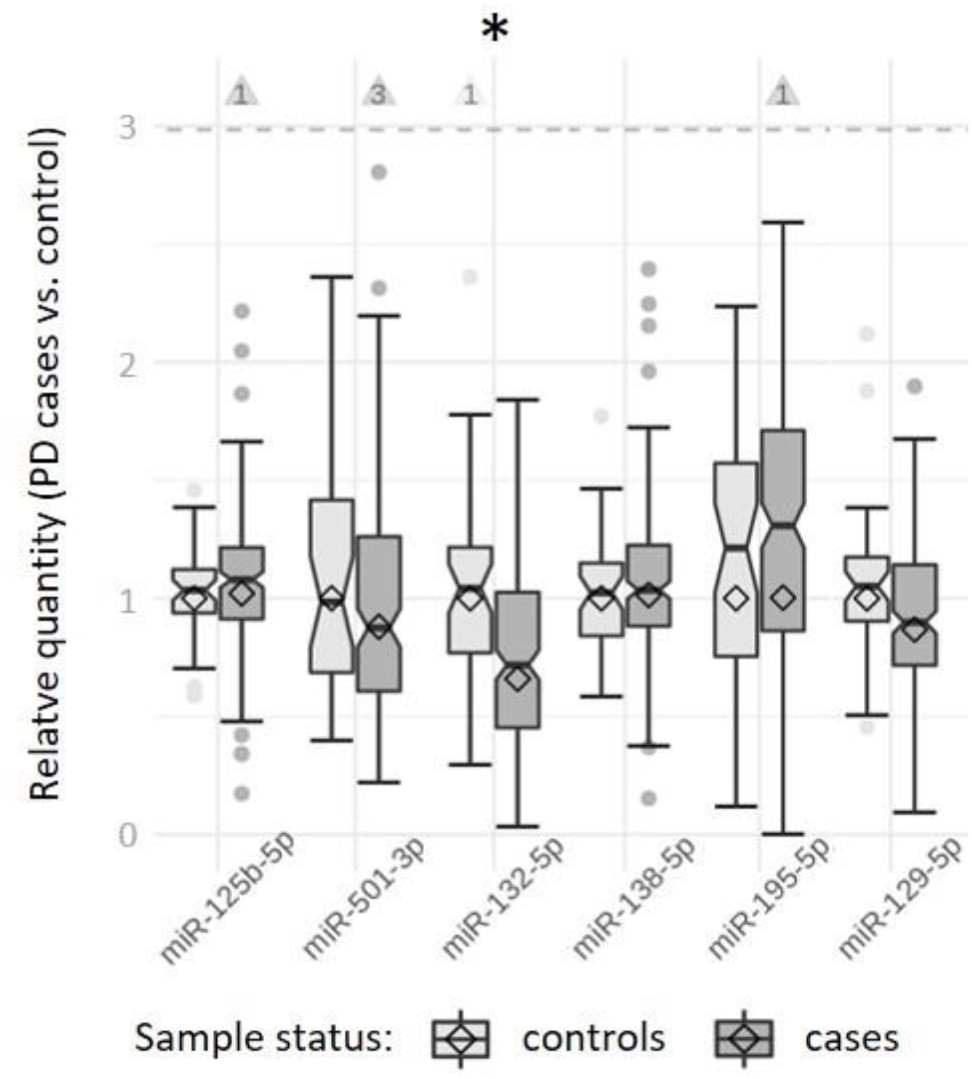
Expression levels of AD candidate miRNAs in *post-mortem* brain samples of PD patients versus controls. <u>Legend</u>. This box and whisker plot displays the relative quantity of Parkinson’s disease (PD) candidate miRNAs in *post-mortem* brain samples of the supratemporal gyrus of 214 Parkinson’s disease (PD) patients vs. 47 controls. The relative quantity of miRNA expression was calculated using the ΔΔCt method. * = nominally statistically significant difference (α=0.05). Also see legend to Figure 2 for more details.

### Meta-analysis of novel miRNA expression results with published evidence

To assess the overall evidence for differential expression of the ten PD and AD candidate miRNAs tested in PD in this study, we added our novel molecular results to the meta-analyses of our previous systematic review (8), resulting in a substantial increase of the sample sizesavailable for the respective miRNAs, by ^~^4-fold on average. At the same time, we added the results from two new smaller studies (19,20), which were published since our initial data freeze. This increase in available data allowed us to meta-analyze two AD candidate miRNAs (hsa-miR-132-5p and hsa-miR-195-5p), for the first time in PD (**Table 3**). Previously, these two miRNAs lacked sufficient data for meta-analysis (8).

**Table 3.**
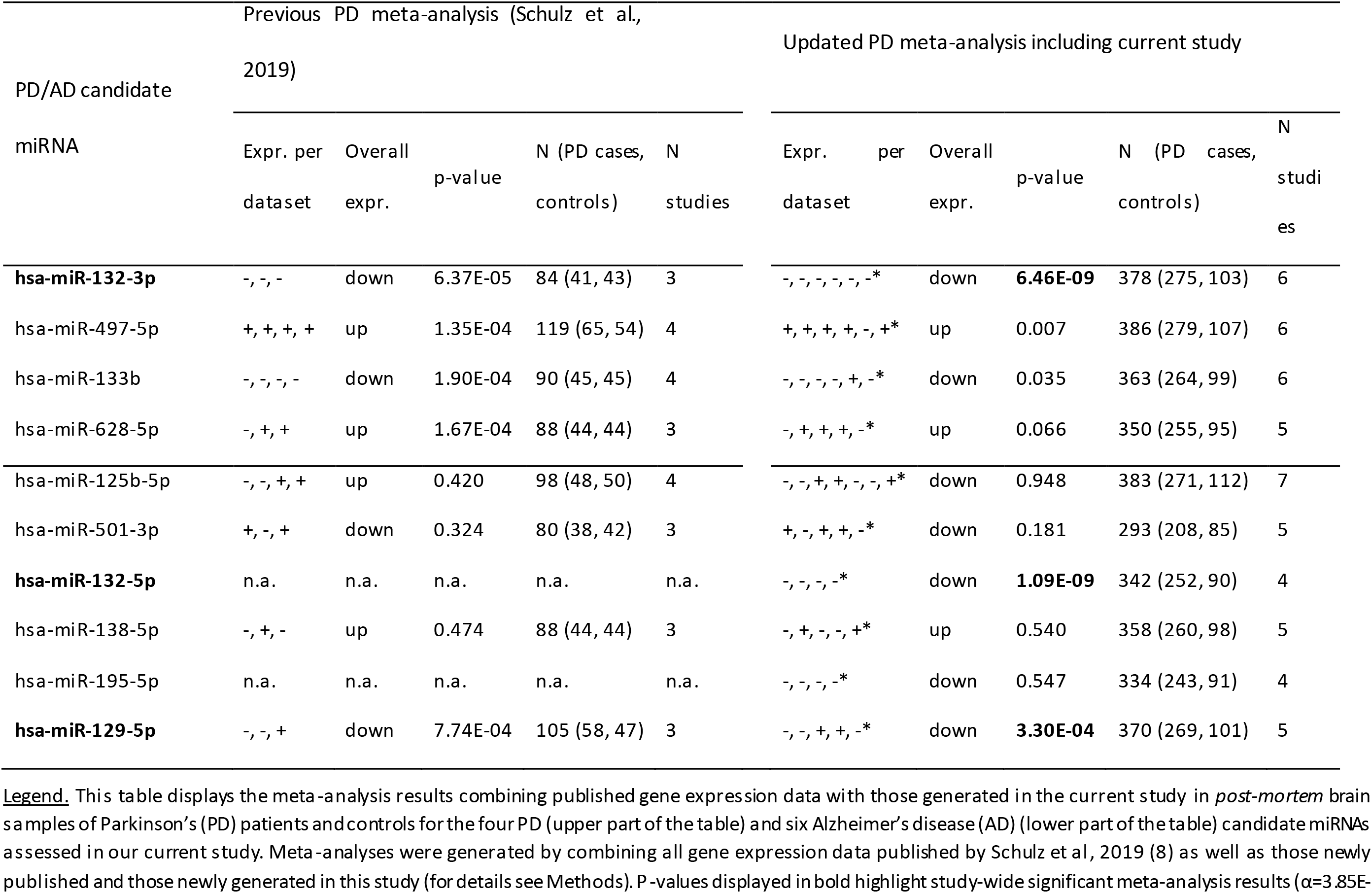
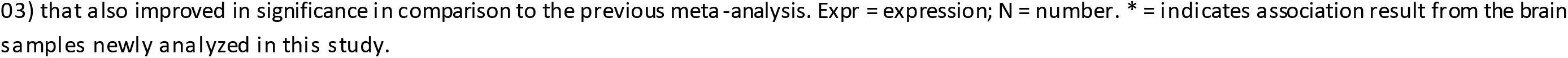
Summary of meta-analysis results for PD- and AD-candidate miRNAs in PD brain samples

The updated meta-analyses showed that hsa-miR-132-3p is significantly differentially expressed in PD vs control brains (**Table 3**). The statistical support of this association is much stronger now than in our previous study (p = 6.46E-09 compared to p = 6.37E-05, respectively, **Table 3**). Second, as expected, the remaining three PD candidate miRNAs, which were not validated in our novel PD brain dataset, no longer reached study-wide significance (α = 3.85E-03) in the updated meta-analyses (**Table 3**). Third, meta-analyses of the six AD candidate miRNAs revealed novel and study-wide significant differential expression for hsa-mir-132-5p (p = 1.09E-09) and for hsa-mir-129-5p in PD brains (p = 3.30E-04, **Table 3**). Finally, we observed a novel finding of study-wide significant effects for hsa-miR-132-3p (1.69E-17 compared to p = 0.03 in the previous meta-analysis (9)) but not for the three other PD candidate miRNAs (hsa-miR-133b, hsa-miR-497-5p, hsa-miR-628-5p; **Table 4**).

**Table 4.**
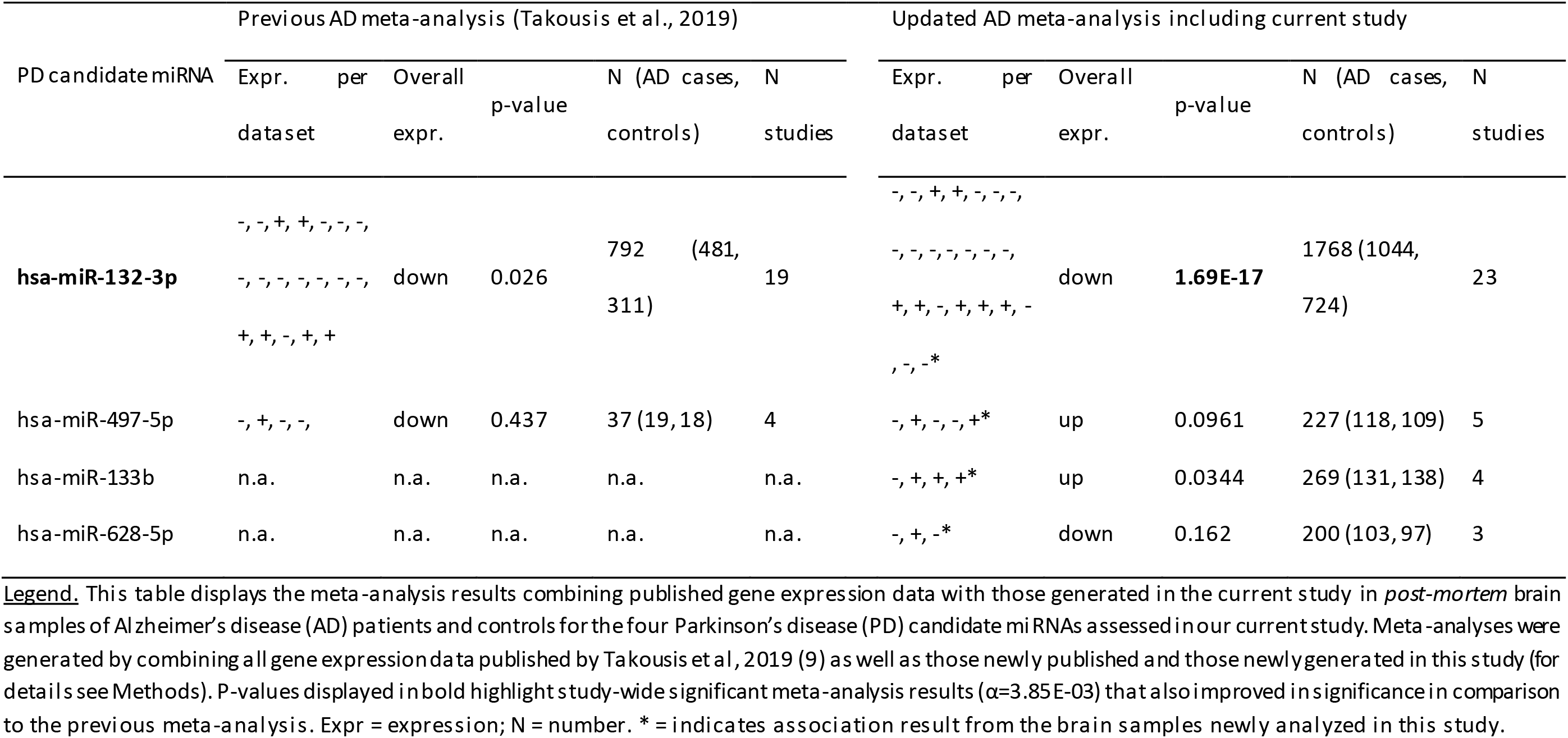
Summary of meta-analysis results for PD candidate miRNAs in AD brain samples

### Linear regression analyses of hsa-miR-132-3p/-5p and hsa-miR-129-5p on PD neuropathology

To gauge at which phase in the disease trajectory the two new PD-linked miRNAs hsa-miR-132-3p/-5p and hsa-miR-129-5p show the strongest degree of differential expression, we assessed their association with α-synuclein and tau Braak staging in the brains of PD patients (for the distribution of Braak and tau staging in the PD brains, see **Additional file 1, Table S1**). Interestingly, hsa-miR-132-3p/-5p expression showed statistically significant association (α = 0.0125) with α-synuclein staging (p = 3.51E-03 for hsa-miR-132-3p, p = 0.0117 for hsa-miR-132-5p) but no or only nominally statistically significant association with tau staging (p = 0.119 and p = 0.0235, respectively; **Figure 4, Additional file 4, Table S3**). In contrast, hsa-miR-129-5p did not show significant association with either neuropathological staging parameter (**Additional file 4, Table S3**).

**Figure 4.**
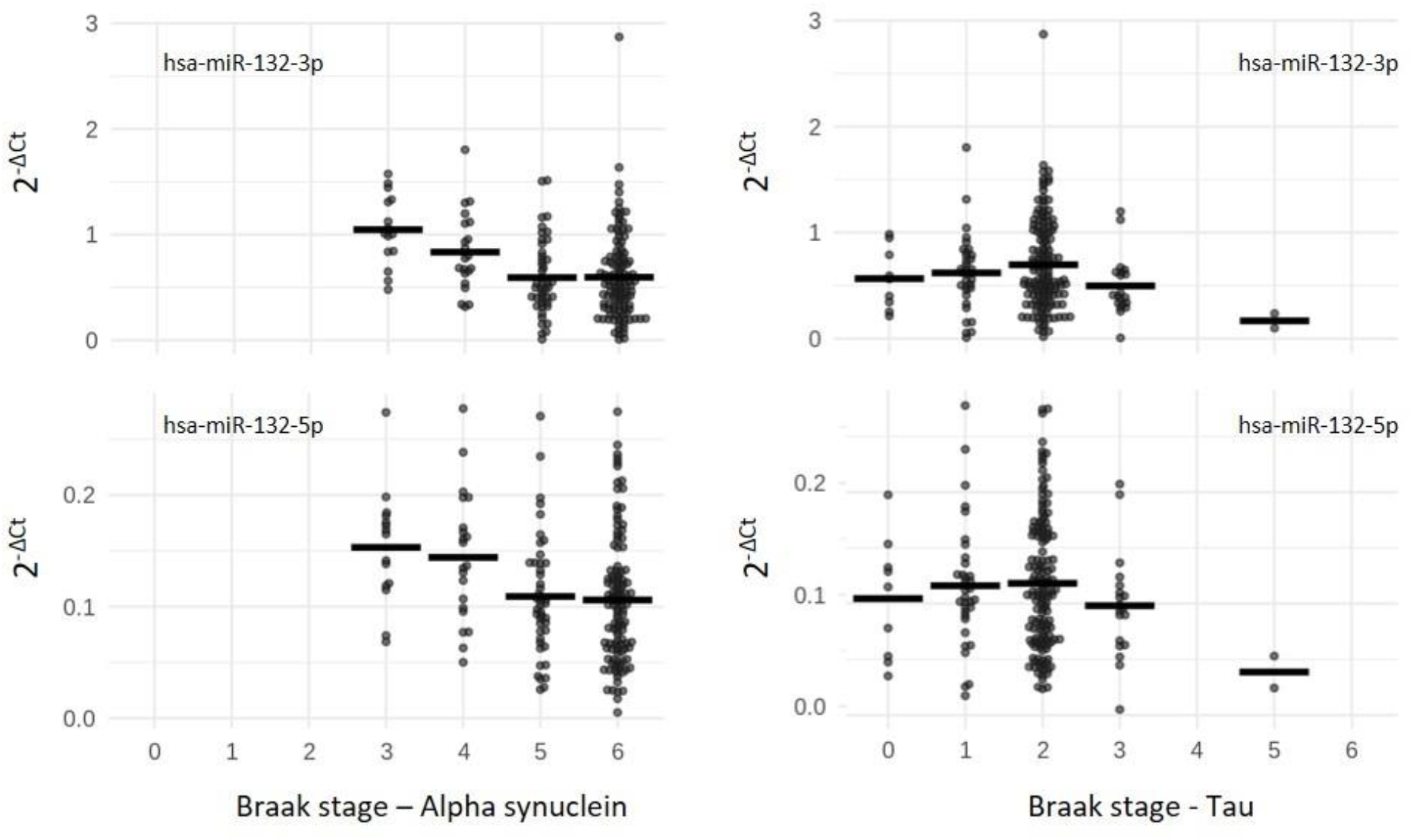
Expression of hsa-miR-132-3p/-5p in PD brains dependent on the neuropathological α-synuclein and tau Braak staging. <u>Legend</u>. This plots display the gene expression of hsa-miR-132-3p and hsa-miR-132-5p normalized to endogenous control assay (dCt) dependent on α-synuclein and tau Braak staging in *post-mortem* brain samples of 213 PD patients. Horizontal lines indicate the mean of the group.

## Discussion

Our study represents the first independent assessment of ten miRNAs showing prior evidence of differential expression in *post-mortem* brain samples in previous work from our group for PD (8,9) and AD (9). To this end, we collected and analyzed one of the largest case-control collections of PD and AD *post-mortem* brain samples (total n = 458) available in the field. Our study provides compelling evidence that hsa-miR-132-3p/-5p and hsa-miR-129-5p show significant differential expression in both PD and AD, potentially indicating shared pathogenic mechanisms across both these neurodegenerative diseases. Furthermore, hsa-miR-132-3p/-5p showed association with neuropathological α-synuclein Braak staging in PD cases, suggesting that it may play a role in α-synuclein aggregation, along the progressive disease course of PD. Intriguingly, hsa-miR-132 directly binds to SNCA mRNA as evidenced by next-generation sequencing (NGS)-based HITS-CLIP data in human brain (21) possibly pinpointing to novel therapeutic approaches in fighting Parkinson’s disease.

Prior evidence for differential gene expression of hsa-miR-132-3p/-5p and hsa-miR-129-5p in *post-mortem* PD vs control brains was very sparse as only few studies using limited sample sizes (ranging from 9 to 62) had reported data on these miRNAs (22–24). Accordingly, our previous meta-analyses (8) were only based on a total of 84 (hsa-miR-132-3p) and 105 (hsa-miR-129-5p) brain samples, while hsa-miR-132-5p could not be meta-analyzed due to a lack of sufficient data. The data from this study - comprising 261 *post-mortem* brain samples of PD and control individuals - quadruplicated the sample sizes available for these miRNAs. Therefore, for the first time, this study now allows to generate robust and reliable results pinpointing to a pivotal role of hsa-miR-132-3p/-5p and hsa-miR-129-5p in the course of PD.

There are several potential limitations of our study. Firstly, while our dataset represents one of the largest collection of *post-mortem* case-control brain samples in PD, our sample size may still be underpowered to detect subtle differences in miRNA expression. As a result, some of the non-validations of the candidate miRNAs assessed in this study may represent false negative findings. Secondly, similar to almost all other recent gene expression studies in the field, we used bulk tissue to quantify miRNA expression. Thus, the reported differential miRNA expression results may at least in part be the result of differences in cell-type composition of our sample rather than a *bona fide* downregulation of the miRNA on a cellular level. This limitation might become addressable by using single-cell/-nucleus sequencing, which was beyond the scope of this work. Thirdly, as our study compared gene expression in *post-mortem* tissue of diseased individuals and unaffected controls, we cannot reliably distinguish cause-effect relationships regarding the role of hsa-miR-132-3p/-5p and hsa-miR-129-5p in PD (or AD). Interestingly, the analyses using neuropathological α-synuclein Braak staging in PD cases suggest that hsa-miR-132-3p/-5p may play a role in the progressive accumulation of α-synuclein along the disease course, beyond the initial disease stages. However, this does not exclude its involvement in pathomechanisms in early disease phases. In contrast, miRNA hsa-miR-129-5p did not show significant association with neuropathological staging in PD cases, possibly suggesting a role predominately at earlier disease stages.

Despite these limitations, we note that several lines of independent evidence support a role of both miR-132 and miR-129-5p in PD and AD: For instance, in PD, downregulation of the miR-132-3p/miR-212-3p cluster (the precursor hsa-mir-132 and precursor hsa-mir-212 share the same primary transcript (25)) was reported to occur in α-synuclein (A30P)-transgenic mice (26). Secondly, it has been shown that deleting the miR-132/212 cluster in mouse models led to impaired memory (27) and increased Aβ production as well as amyloid plaque formation (28). Third, both hsa-miR-132-3p and miR-129-5p emerged among miRNAs with neuroprotective roles against amyloid β-peptide accumulation in primary mouse and human neuronal cell culture models (29). Lastly, a neuroprotective role for miR-129-5p has recently been reported in a rat model of AD (established by injecting Aβ25-35 into the brain), where miR-129-5p inhibited neuronal apoptosis (30).

## Conclusions

In conclusion, our study provides novel and compelling evidence that hsa-miR-132-3p/-5p and hsa-miR-129-5p are differentially expressed in *post-mortem* brain samples of both PD and AD. Future work needs to elucidate whether these concurrent miRNA expression results are the result of shared pathogenic mechanisms across both of these diseases and to delineate the precise molecular mechanisms underlying these associations.

## List of abbreviations

AD: Alzheimer’s disease
GLM: generalized linear model
miRNA: micro RNA
PD: Parkinson’s disease
PMI: *post-mortem* interval
QC: quality control
RIN: RNA integrity number
STG: superior temporal gyrus

## Declarations

### Ethics approval and consent to participate

All participants had given prior written informed consent for the brain donation. The tissue bank activities of the Parkinson’s UK Brain Bank were approved by the Ethics Committee for Wales (ref 18/WA/0238), and the Oxford Brain Bank activities were approved by the Ethics Committee of the University of Oxford (ref 15/SC/0639).

### Consent for publication

Not applicable.

### Availability of data and materials

All data generated or analyzed during this study are available from the corresponding author on request.

## Authors’ contributions

1. Research project A. Conception, B. Organization, C. Execution;
2. Statistical Analysis: A. Design, B. Execution, C. Review and Critique;
3. Manuscript Preparation: A. Writing of the first draft, B. Review and Critique.

VD: 1A, 1B, 1C, 2C, 3A

MS: 2B, 3B

IF: 1C, 3B

DG: 1C, 3B

JS: 2B, 3B

LM: 1A, 3B

SG: 1C, 3B

LP: 1C, 2C, 3B

LB: 1A, 1B, 1C, 2C, 3B

CL: 1A, 1B, 2A, 2C, 3A, 3B

## Acknowledgements

The authors are indebted to all tissue donors of the Brain Banks, who have graciously allowed us to conduct this study.

## Additional material

**Additional file 1:** Additional file 1.docx, **Table S1.** Overview of the PD case-control post-mortem brain samples analyzed in this study.

**Additional file 2:** Additional file 2.docx, **Table S2.** Overview of the AD case-control *post-mortem* brain samples analyzed in this study.

**Additional file 3:** Additional file 3.docx, **Figure S1.** Box plot displaying the distribution of qPCR-based Ct values for AD samples analyzed in this study.

**Additional file 4:** Additional file 4.docx, **Table S3.** Linear regression analysis of hsa-miR-132-3p/-5p and hsa-miR-129-5p on α-synuclein and tau Braak staging in *post-mortem* PD brain samples.

