## Additional file 1 for "Common signatures of differential microRNA expression in Parkinson’s and Alzheimer’s disease brains"

**Table S1.** Overview of the PD case-control *post-mortem* brain samples analyzed in this study

| | | PD cases | | PD controls |
|---|---|---|---|---|
| **Total number** | | 214 | | 47 |
| **Sex** | | | | |
| | Number of males (%) | 144 (67.3%) | | 26 (55.3%) |
| | p-value[a] | | 0.145 | |
| **Age at death (years)** | | | | |
| | average (±SD) | 78.8 (6.7) | | 81.1 (9.7) |
| | median (IQR) | 79 (75-84) | | 83 (75.5-89) |
| | range | 57-93 | | 58-96 |
| | p-value[b] | | 0.137 | |
| **Age at onset (years)**[c] | | | | |
| | average (±SD) | 65.6 (10.2) | | n.a. |
| | median (IQR) | 67 (59-72) | | n.a. |
| | range | 30-88 | | n.a. |
| **Disease duration (years)**[c] | | | | |
| | average (±SD) | 13.0 (7.3) | | n.a. |
| | median (IQR) | 11 (8-17) | | n.a. |
| | range | 1-44 | | n.a. |
| **PMI (hours)** | | | | |
| | average (±SD) | 19.4 (9.5) | | 21.1 (9.7) |
| | median (IQR) | 19 (12-24) | | 21 (13.5-24.75) |
| | range | 2-48 | | 5-48 |
| | p-value[b] | | 0.302 | |
| **RIN value** | | | | |
| | average (±SD) | 3.7 (1.4) | | 3.6 (1.4) |
| | median (IQR) | 3.05 (2.5-4.68) | | 2.8 (2.4-4.8) |
| | range | 2.2-7.5 | | 2.3-7.6 |
| | p-value[b, d] | | 0.629 | |
| **RNA A260/280** | | | | |
| | average (±SD) | 1.94 (0.03) | | 1.95 (0.05) |
| | median (IQR) | 1.94 (1.92-1.95) | | 1.94 (1.93-1.96) |
| | range | 1.87-2.06 | | 1.89-2.20 |
| | p-value[b] | | 0.130 | |
| **Alpha-synuclein Braak stage**[e] | | | | |
| | Stage 3 | 15 (7.0%) | | n.a. |
| | Stage 4 | 21 (9.8%) | | n.a. |
| | Stage 5 | 46 (21.5%) | | n.a. |
| | Stage 6 | 120 (56.1%) | | n.a. |
| | Data unavailable | 12 (5.6%) | | n.a. |
| **Tau Braak stage**[f] | | | | n.a. |
| | Stage 0 | 9 (4.2%) | | n.a. |
| | Stage 1 | 35 (16.5%) | | n.a. |
| | Stage 2 | 146 (68.2%) | | n.a. |
| | Stage 3 | 20 (9.3%) | | n.a. |
| | Stage 5 | 2 (0.9%) | | n.a. |
| | Data unavailable | 2 (0.9%) | | n.a. |

Legend. *Post-mortem* brain samples from the supratemporal gyrus of Parkinson's (PD) patients and corresponding controls were provided by the Parkinson's UK Brain Bank at Imperial College London. SD = standard deviation; IQR = interquartile range; PMI = *post-mortem* interval; RIN = RNA integrity number; [a] = Pearson's Chi-squared test with Yates' continuity correction; [b] = Welch's two-sample t-test; [c] = data available for 155 individuals; [d] = due to some deviation from normality, we also performed Wilcoxon ranksum test, but results did not change substantially, thus only the results of Welch's two-sample t-test are provided here; [e] = data available for 202 individuals; [f] = data available for 212 individuals; n.a. = not applicable/not available.
