## Additional file 2 for "Common signatures of differential microRNA expression in Parkinson’s and Alzheimer’s disease brains"

**Table S2.** Overview of the AD case-control *post-mortem* brain samples analyzed in this study

| | AD cases | AD controls |
|---|---|---|
| **Total number** | 99 | 91 |
| **Sex** | | |
| Number of males (%) | 50 (51%) | 51 (56%) |
| p-value[a] | 0.536 | |
| **Age at death (years)** | | |
| average (±SD) | 81.6 (8.0) | 77.5 (13.8) |
| median (IQR) | 83 (77.5-87) | 81(68.5-88.5) |
| range | 61-95 | 41-100 |
| p-value[b] | 0.0142 | |
| **PMI (hours)** | | |
| average (±SD) | 57.1 (30.6) | 48.41 (31) |
| median (IQR) | 48 (30-73.75) | 48 (24-48) |
| range | 9-140 | 5-168 |
| p-value[b] | 0.0540 | |
| **RIN value** | | |
| average (±SD) | 3.0 (1.2) | 4.2 (1.4) |
| median (IQR) | 2.6 (2.3-3.25) | 4.00 (2.95-5.2) |
| range | 1.2-7.8 | 2.1-7.6 |
| p-value[b, c] | 6.92e-09 | |
| **RNA A260/280** | | |
| average (±SD) | 1.90 (0.04) | 1.93 (0.04) |
| median (IQR) | 1.89 (1.86-1.94) | 1.93 (1.91-1.95) |
| range | 1.80 - 1.99 | 1.80-2.00 |
| p-value[b, c] | 2.063e-05 | |
| **Braak Stage** | | |
| Stage 0 | 0 (0%) | 6 (6.6%) |
| Stage I/II | 0 (0%) | 72 (79.1%) |
| Stage III | 0 (0%) | 6 (6.6%) |
| Stage IV | 8 (8.1%) | 0 (0%) |
| Stage V/VI | 91 (91.9%) | 0 (0%) |
| Data unavailable | 0 (0%) | 7 (7.7%) |

Legend. *Post-mortem* brain samples from the supratemporal gyrus of Alzheimer's (AD) patients and corresponding controls originate from the biobank of the longitudinal, prospective Oxford Project to Investigate Memory and Aging (OPTIMA) at University of Oxford (also described in Dobricic et al., 2021 (Dobricic et al., 2021)). SD = standard deviation; IQR = interquartile range; PMI = *post-mortem* interval; RIN = RNA integrity number; [a] = Pearson's Chi-squared test with Yates' continuity correction; [b] = Welch's two-sample t-test; [c] = due to some deviation from normality, we also performed Wilcoxon rank sum test, but results did not change substantially, thus only the results of Welch's two-sample t-test are provided here; n.a. = not available.
