## Additional file 3 for "Common signatures of differential microRNA expression in Parkinson’s and Alzheimer’s disease brains"

**Figure S1.** Box plot displaying the distribution of qPCR-based Ct values for AD samples analyzed in this study

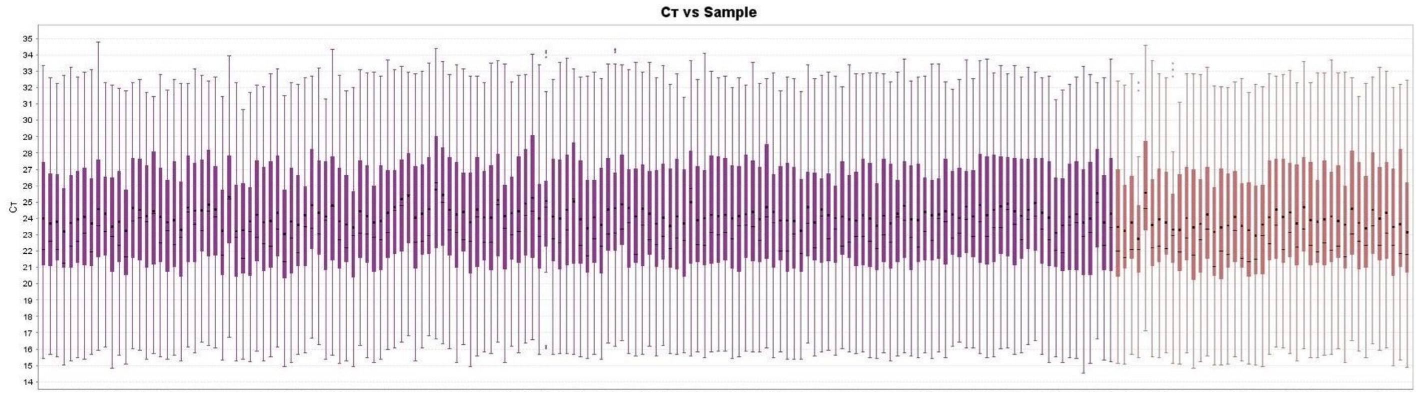


Legend. This figures shows similar distributions of Ct values in samples with lower (RIN <5, represented in purple color) vs higher RIN values (RIN ≥ 5, represented in brown color) values. The horizontal black bar shows the median Ct value. The black circle shows the mean Ct value. The solid box shows the range of the middle 50% of the Ct values for each sample. Note that this figures has already been published in Dobricic et al., 2021 (Dobricic et al., 2021)).
