## Additional file 4 for "Common signatures of differential microRNA expression in Parkinson’s and Alzheimer’s disease brains"

**Table S3.** Linear regression analysis of hsa-miR-132-3p/-5p and hsa-miR-129-5p on α-synuclein and tau Braak staging in *post-mortem* PD brain samples

| MiRNA | Braak staging | Effect estimate (95% CI) | p |
|---|---|---|---|
| **hsa-miR-132-3p** | **α-synuclein** | **-0.211 (-0.352, -0.071)** | **3.51E-03*** |
| | tau | -0.113 (-0.255, 0.028) | 0.119 |
| **hsa-miR-132-5p** | **α-synuclein** | **-0.185 (-0.327, -0.043)** | **0.0117*** |
| | **tau** | **-0.166 (-0.309, -0.024)** | **0.0235** |
| hsa-miR-129-5p | α-synuclein | -0.077 (-0.230, 0.077) | 0.329 |
| | tau | -0.060 (-0.220, 0.100) | 0.463 |

Legend. This table displays the linear regression results of hsa-miR-132-3p/-5p and hsa-miR-1129-3p/-5p on α-synuclein and tau Braak staging in *post-mortem* brain samples of 202 (α-synuclein) and 212 (tau) PD patients. P-values displayed in bold are at least nominally significant, * denotes p-values showing significant results after Bonferroni correction (α=0.0125), also see Methods.
